# The Influence of Aging and Mechanical Stretch in Alveolar Epithelium ER Stress and Inflammation

**DOI:** 10.1101/157677

**Authors:** MS Valentine, JA Herbert, PA Link, F Kamga Gninzeko, MB Schneck, K Shankar, J Nkwocha, AM Reynolds, RL Heise

## Abstract

Ventilator-Induced lung injury (VILI) is a form of acute lung injury that is initiated or exacerbated by mechanical ventilation. The aging lung is more susceptible to lung injury. Harmful mechanical stretch of the alveolar epithelium is a recognized mechanism of VILI, yet little is known about how mechanical stretch affects aged epithelial cells. An activated response known as Endoplasmic Reticulum (ER) Stress occurs at the cellular level, which is increased with aging. The disrupted ER function results in disruption in cellular homeostasis, apoptosis, and inflammation. We hypothesized that age and mechanical stretch increase proinflammatory gene expression that is mediated by ER stress. Type II alveolar epithelial cells (ATII) were harvested from C57Bl6/J mice 8 weeks (young) and 20 months (old) of age. The cells were cyclically mechanically stretched at 15% change in surface area for up to 24 hours. Prior to stretch, groups were administered 4-PBA or vehicle as a control. Mechanical stretch upregulated both ER stress and proinflammatory gene expression in ATIIs. Age-matched and mis-matched monocyte recruitment by ATII conditioned media was quantified. Administration of 4-PBA attenuated both the ER stress and proinflammatory increases from stretch and/or age and significantly reduced monocyte migration to ATII conditioned media. Age increases susceptibility to stretch-induced ER stress and downstream inflammation in a primary ATII epithelial cell model.

## Introduction

Mechanical ventilation frequently exacerbates underlying pulmonary conditions and produces an exaggerated inflammatory response that potentially leads to sepsis and multisystem organ failure (40). This exacerbation or injury is classified as Ventilator-Induced Lung Injury (VILI). The pathophysiology of VILI is characterized by an exaggerated proinflammatory cytokine release and influx of inflammatory cells, loss of alveolar barrier integrity and subsequent pulmonary edema formation, decreased lung compliance, and profound hypoxia. These features reflect three integrated mechanisms of injury: alveolar over-distention, cyclic atelectasis, and inflammatory cell activation (19, 46). These physical injury mechanisms are frequently modeled *in vitro* with mechanical cyclic stretch using lung epithelial or endothelial cells.

Alveolar epithelial cells contribute to the initiation, amplification, and resolution of inflammatory processes in the lung. Epithelial cells produce inflammatory mediators assumed to be involved in the recruitment and regulation of macrophages (13, 55). Furthermore, cyclic stretch on epithelial cells grown on deformable membranes has also shown to cause injury and induce cytokine release by alveolar epithelial cells (58). Cyclic stretch of alveolar epithelial cells resulted in increased cell injury and death, increased apoptosis, increased acidification and bacterial growth, and increased general inflammatory response, often shown with amplified gene expression and release of IL-6 and IL-8 (11, 29, 58). Together, these observations validate that alveolar epithelial cells are involved with the mechanotransduction and exaggerated inflammatory response that is characteristic of VILI.

Mechanical ventilation leads to poorer outcomes in the elderly population. Mortality rates and hospital discharge to extended care facilities increased consistently for each decade of age over the age of 65 years in mechanically ventilated patients (20). Epidemiological studies suggest that age is a predictive factor in the severity of VILI; however, the exact molecular mechanisms between age and VILI are still unknown (14, 47, 51). In rodent models of VILI, we and others have shown that age increases susceptibility to ventilator-induced edema, injury, and mortality (1, 31). In general, advanced age is also known to promote an increasingly dysregulated innate immune/inflammatory response to injury with an overall shift towards a proinflammatory state (21). Aging promotes chronic inflammation in the murine lung (2) and inhibits repair of the lung epithelium following influenza insult (62). Better understanding these age-associated interactions are critical to developing therapeutic approaches that aim to target the possible age-associated mechanisms of injury related to the pathophysiology of VILI.

One potential regulator of age-associated inflammation is the endoplasmic reticulum (ER). The ER is a multifunctional organelle responsible for lipid biosynthesis, calcium storage, and protein folding and processing. Due to the innate inefficiency of protein folding, there can be as many as 30% of proteins that never acquire their fully folded, programmed conformation (49). Prolonged unresolved ER stress can lead to the accumulation of misfolded proteins, disruption of cellular functions, cellular apoptosis, and has been shown to play a key role in many chronic inflammatory disease states (17). Specifically, ER stress has been shown to regulate apoptosis and epithelial to mesenchymal transition in alveolar epithelial cells (63).

ER-stress also becomes increasingly dysregulated with age (45). There is a general age-associated increase in the occurrence of protein misfolding and accumulation. Unsurprisingly ER stress is implicated as a promoter of many pathological disease states associated with aging (6, 45). Additionally, lung-related ER stress is implicated in the age-associated increase in pulmonary fibrosis (36).

While numerous studies have shown convincing evidence for the dominant role of ER stress in various inflammatory disease states, there have only been a few studies investigating ER stress in the context of acute lung injury and/or ventilator-induced lung injury. Several ER stress pathway proteins are key modulators of epithelial permeability and barrier dysfunction in young mice and rats (17, 63). It was demonstrated that extended epithelial stretch activates ER stress pathways, which resulted in increased alveolar permeability, cell death, and proinflammatory signaling (16); however, these implications have yet to be investigated in an aging model. To further validate that therapeutic targeting of the ER stress response may attenuate VILI, these inferences need to be investigated in the context of aging to understand the potential therapeutic targets.

We hypothesized that the alveolar type II (ATII) cells respond age-dependently to mechanical stretch with increased inflammation and monocyte recruitment, dependent upon ER stress. To investigate this relationship, we isolated and cultured primary alveolar epithelial ATIIs from young and old murine subjects. These cells were exposed to cyclic mechanical stretch to model alveolar distension. We measured age-associated differences in ER stress response, cell injury/inflammation, and apoptosis. We inhibited ER stress using the chaperone 4- phenylbutyrate (4-PBA) (35). Additionally, we quantified the migration of bone marrow-derived monocytes to ATII conditioned media.

## Materials and Methods

All animals are housed in accordance with guidelines from the American Association for Laboratory Animal Care and Research protocols and approved by the Institutional Animal Care Use Committee at Virginia Commonwealth University (Protocol No. AD1000009).

### ATII cell isolation and culture

We harvested, isolated, and cultured ATII primary alveolar epithelial cells from young (8 weeks) and old (20 months) C57BL/6J wild-type mice using previously cited methods (12). ATII were then cultured in Bronchial Epithelial Cell Growth Media (BEGM, Lonza), with the included supplements except for hydrocortisone, supplemented with 10 ng/ml keratinocyte growth factor (KGF, PeproTech). ATII were found to be greater than 93% pure in both young and old cultures by staining for positive pro-surfactant C and the inclusion of lamellar bodies. For stretch experiments, cells were plated onto Collagen I-coated, 6-well silicone bottomed plates (Flexcell International Corp., BF-3001A BioFlex) and cultured for 48 hours prior to mechanical stimulation.

### Cell Stretch

Using the Flexcell Tension Plus System (Flexcell Inc), we applied radial, cyclic (0.86Hz) stretch corresponding to a 15% change in surface area. Statically cultured ATII cells were used as controls. Cells underwent stretch or static conditions, and after 4 hours or 24 hours, RLT buffer (Qiagen) or 4% paraformaldehyde were added to wells for further processing. Cell supernatants were collected for cytokine analysis and conditioned media experiments.

### ER Stress Intervention

One hour before mechanical stretch, each well received either 20ul vehicle (ultrapure water) or 10 mM sodium 4-phenylbultyrate (PBA) (Calbiochem, San Diego, CA) in ultrapure water. 4-PBA is already in use clinically as a successful treatment for systemic inflammatory diseases such as diabetes (61).

### Immunofluorescence Staining for CHOP, ATF4

Fixed wells were probed for CHOP (L63F7) mouse mAB (1:3200) and ATF-4 (D4B8) rabbit mAB (1:200) using secondary antibodies anti-mouse Alexa Fluor® 594 conjugate (1:250) and anti-rabbit Alexa Fluor® 488 conjugate respectively. All primary and secondary antibodies were obtained from Cell Signaling Technology (Danvers, MA, USA). The counterstaining was done using Prolong® Gold antifade mounting with DAPI (ThermoFisher, Waltham, MA, USA). Finally, samples were imaged with an Olympus IX71 under appropriate emission/excitation wavelengths.

### Quantitative real-time Polymerase Chain Reaction

Total RNA was isolated and purified from each treatment group using RLT buffer (Qiagen) and the RNeasy mini kit (Qiagen, Valencia, CA). We then synthesized the complementary DNA using the iScript RT kit (Biorad). For cDNA from the ATII primary alveolar epithelial cell, we used custom QPCR plates (Biorad) to perform an analysis of inflammatory genes of interest. The genes included were the following: Ccl12, Ccl2, Ccl20, Ccl3, Ccl4, Ccl6, Ccl7, Ccl9, Ccr1, Csf1, Il10rb, Il15, Il1a, Il1r1, Il2rg, Il6st, Sgpp1, Tnfrsf11b, and Vegfa. Additional primers for analysis of ER stress-related genes, CHOP and ATF4, were purchased from Integrated DNA Technologies. QPCR was performed using Sybr Green (Applied Biosystems) and the CFX96 Touch^TM^ Real-Time PCR Detection System (Biorad). Data was analyzed using the 2^-ΔΔCT^ method, and target genes were normalized to two housekeeping genes using ribosomal 18s and GAPDH.

### Inflammatory Mediator Analysis

We measured the concentrations of MCP-1 (CCL2) and MIP-1β (CCL4) inflammatory cytokines in the collected cell media of each experimental group using MCP-1 (DY479) and MIP-1β (DY451) Mouse DuoSet ELISA kits (R&D Systems) according to the manufacturer’s instructions.

### Monocyte Invasion Assay

Bone marrow derived monocytes (BMDMs) were isolated from young (8 weeks) and old (20 months) C57BL/6J mice, as described by Trouplin et al (57). Monocyte migration was then evaluated using an invasion assay, performed as described by Murray et al (44), with minor modifications. BMDMs were seeded at a density of 1 × 10^5^ cells/100ul BMDM growth media without FBS, on Collagen I-coated (Sigma-Aldrich) Transwell inserts with 8.0 um pore sizes (Corning, USA). 0.6 ml of BEGM or conditioned media from the ATII 24 hour groups were placed in the reservoir.

A Live/Dead Viability Assay (ThermoFisher) was used to quantify cell invasion through the Transwell membrane. Live/Dead images were taken immediately after the staining procedure an Olympus IX71 Microscope (Olympus). Total cell counts were performed using ImageJ’s particle analysis function with the following inclusion parameters: Size (in Pixels): 10-120. Circularity: 0.10-0.99.

## Statistics

A total of 114 young and old mice were used for this study. All experiments were performed with a minimum of n=3 primary cell isolations in triplicate wells. Larger n values were utilized where possible. Limitations exist in the number of 20-month-old mice available from the National Institute on Aging. Therefore minimum numbers to achieve statistical significance via a power analysis were utilized. Results are presented as mean +/- SD. GraphPad Prism was used for all statistical analyses. For multiple-group comparisons, we used a two-way analysis of variance (ANOVA) with age and stretch as factors, followed by a post-hoc Tukey test to determine significance. P<0.05 was considered statistically significant.

## Results

We quantified inflammatory gene expression involved in leukocyte recruitment and injury signaling. We used these inflammatory arrays to compare the gene expression profile of mechanically stretched alveolar epithelial type II (ATII) cells to that of static controls. Fold changes of gene expression for ATII cells stretched or static for 4 hours are shown in Figure 1 for the following genes: Monocyte Chemoattractant Protein 1 (MCP-1/CCL2), Macrophage Inflammatory Protein 1 alpha (MIP-1α/CCL3), Macrophage Inflammatory Protein 1 beta (MIP-1β/CCL4), Monocyte Chemotactic Protein 3 (MCP-3/CCL7), Monocyte Chemotactic Protein 5 (MCP-5/CCL12), Macrophage Inflammatory 3 alpha (MIP-3α/CCL20), and Interleukin 6 (IL-6). Fold changes in gene expression are compared to the Young AT2 static group. Age and stretch resulted in significant increases in gene expression for MCP-1, MIP-1α, and MCP-5. MCP-3 showed significantly increased gene expression between Young and Old static groups and between Young static and Young stretched, but not between Young static and Old Stretch, or Old static and Old stretched groups. MIP-1β and MIP-3α resulted in significantly increased gene expression between Young static and Young stretched cells, Old static and Old stretched cells, and Young and Old stretched cells. However, there was no significant difference in MIP-1β or MIP-3α gene expression between Young and Old static groups. Interleukin 6 signal transducer (IL-6st) activation depends upon the binding of cytokines to the IL-6 receptor, which is involved in a wide variety of inflammatory disease states, including VILI (24). IL-6st was also significantly upregulated in our old static cells compared to young static cells; however, there was no difference with stretch. There were no statistically significant changes in the other genes we examined between any groups; therefore, they are not shown in Figure 1.

**Figure 1:**
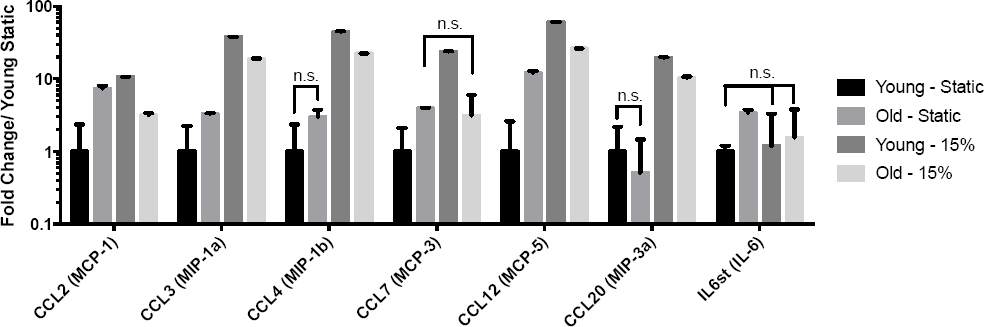
Cyclic Stretch (15%) +/- Age Upregulate Inflammatory Gene Expression in ATII cells at 4 hours. Columns are fold change differences in gene expression compared with Young Static placed on a log10 scale to observe large changes in expression. Data are presented as mean +/- st. deviation, n ≥ 3 per group. Unless noted with n.s., representing no significance, each group was significantly different, p<0.05, compared with all other groups.

After quantifying cytokine gene expression, we examined ER stress as a potential regulator. In order to evaluate alveolar epithelial ER stress responses of cells exposed to mechanical stretch, we investigated the expression of two key indicators of the ER stress pathway: Activating Transcription Factor 4 (ATF4) and C/EBP homologous protein expression (CHOP) (Fig 2). We compared the young and old gene expression of both CHOP and ATF4 between stretched and statically cultured cells. We observed significantly increased CHOP in response to advanced age alone, mechanical stretch alone, and in combination. CHOP is a transcription factor activated by ER stress, which is believed to help mediate cellular apoptosis and inflammation (17). Interestingly, we did not see the same age‐ and stretch-induced upregulation with ATF4, which is another known marker for the ER stress response. However, we observed significantly increased ATF4 expression in the old cells in response to mechanical stretch. After administrating 4-PBA, we observed significant decreases in the ER stress marker CHOP in young and old cells that received mechanical stretch and in statically cultured old cells; however, CHOP expression did not change in statically cultured young cells (Fig 2). 4-PBA administration also attenuated increased ATF4 expression in the old stretched cells; however, there was no change in young stretched cells, or in the statically cultured cells.

**Figure 2:**
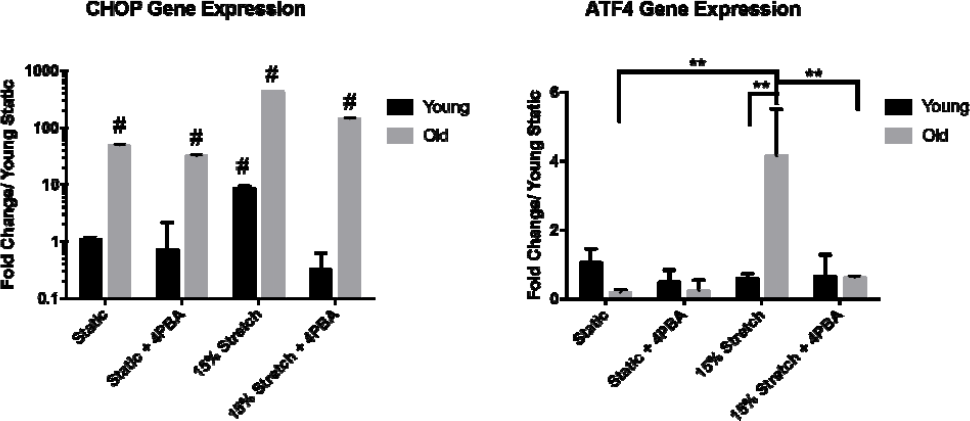
(A) CHOP and (B) ATF4 gene expression after 24 hours of stretch or static culture +/- 4-PBA. Columns are fold change differences in gene expression to young static1. CHOP is shown on a log10 scale to observe large changes in gene expression. Data are presented as mean +/- st. deviation, n ≥ 3 per group. (A) # p<0.0001 compared with all other groups with symbol, (B) ** = p<0.01, as indicated.

The same trends in ATF4 and CHOP gene expression were observed through immunofluorescent staining. Old ATII cells were more positively stained for CHOP under static conditions when compared to the Young ATIIs (Fig. 3A and 3E). Also, the presence of CHOP is more accentuated with stretch in the Young ATIIs (Fig. 3A and 3C). Additionally, 4-PBA attenuated CHOP (Fig. 3B, 3D, 3F, and 3H) compared to vehicle controls (Fig. 3A, 3C, 3E, and 3G). Similarly, Old ATII cells were more positively stained for ATF4 under static conditions when compared to the young (Fig. 3I and 3M), though the difference is less pronounced compared to the CHOP stained images. Similarly, the presence of ATF4 is more accentuated with stretch in the Old ATIIs (Fig. 3M and 3O). 4-PBA also attenuated the positive staining of ATF4 (Fig. 3B, 3D, 3F, and 3H) compared to vehicle controls (Fig. 3A, 3C, 3E, and 3G).

**Figure 3:**
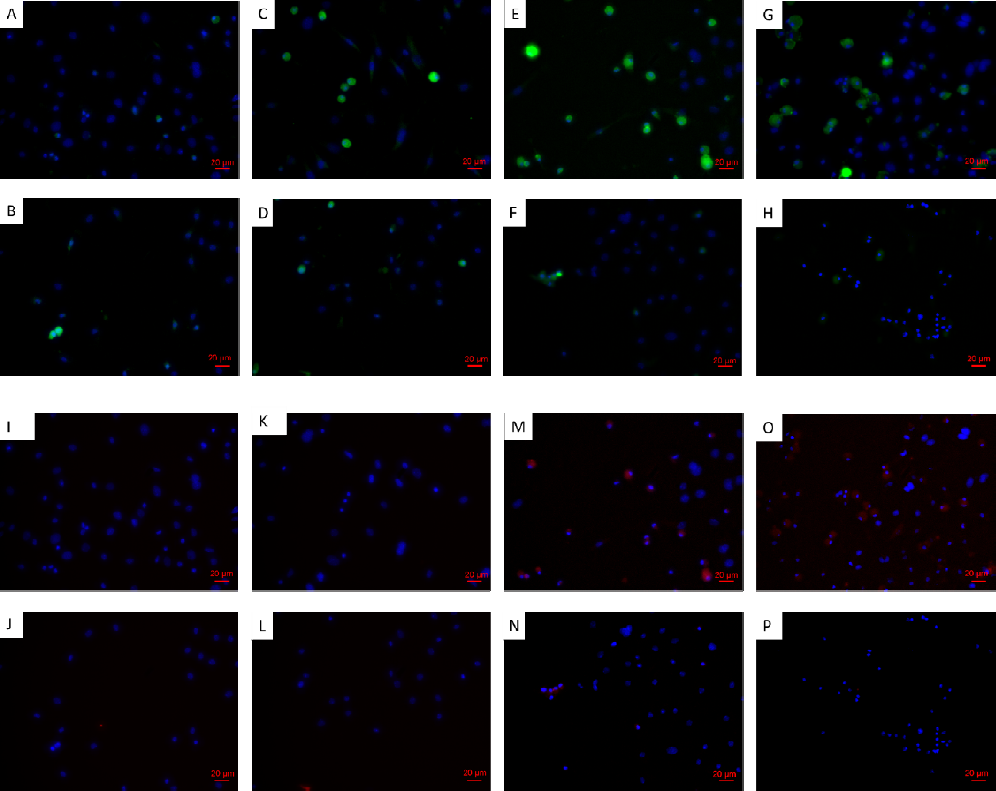
Cyclic Stretch (15%) for 24hrs +/- Age influences detection of (A-H) CHOP and (I-P) ATF4 in ATII Cells. Representative images were chosen from 3 wells stained per each experimental condition: A&I: Young Static, B&J: Young Static + 4PBA, C&K: Young Stretch, D&L: Young Stretch + 4PBA, E&M: Old Static, F&N: Old Static + 4PBA, G&O: Old Stretch, and H&P: Old Stretch + 4PBA. (A-H): DAPI (blue) and CHOP (green). (I-P): DAPI (blue) and ATF4 (red). Scale bars represent 20 μm.

We then examined the influence of 4PBA on two key factors that would influence inflammatory cell recruitment, MCP-1 (CCL2) and MIP-1β (CCL4), which we also saw influenced by age and mechanical stretch (Fig 1). Gene expression for MCP-1 remained significantly elevated after 24 hours when comparing stretched Old ATII cells to static Old ATII cells. Furthermore, MCP-1 (CCL2) gene expression was significantly increased in Old ATII cells to Young ATII cells with mechanical stretch; however, there was no significant difference in gene expression between Old ATII and Young ATII static cells or between Young stretched and static ATII cells (Fig 4A). 4-PBA significantly decreased MCP-1 gene expression in stretched Old ATII cells, but 4-PBA did not significantly decrease MCP-1 gene expression in the stretched Young ATII cells or the young and old static cells (Fig. 4A). Differing from the gene expression at 4 (Fig. 1) or 24 (Fig. 4A) hours, MCP-1 concentration in the media was significantly higher after 24 hours in stretched Young ATII cells compared to static Young ATII cells or stretched Old ATII cells (Fig. 4B). 4-PBA significantly lowered MCP-1 secretion in both static and stretched Young and Old ATII cells (Fig. 4B).

**Figure 4:**
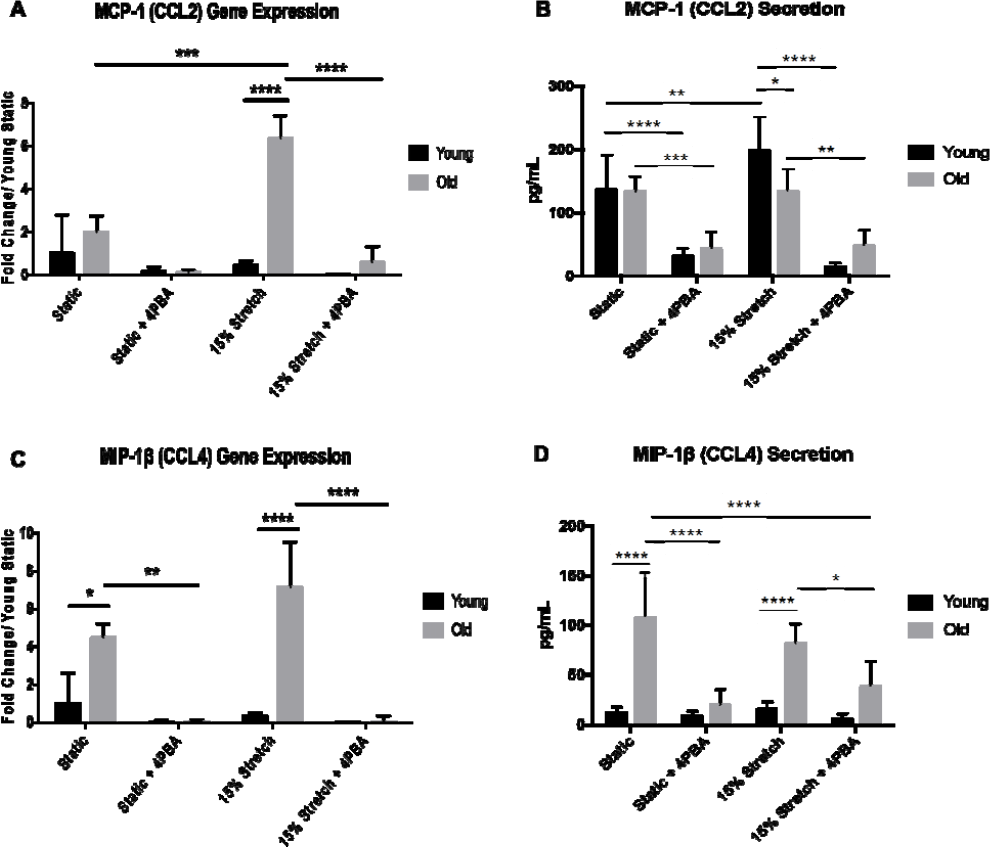
(A) MCP-1 (CCL2) gene expression and (B) cytokine secretion and (C) MIP-1β (CCL4) gene expression (D) and cytokine secretion in ATII cells after 24 hours. Gene expression data are fold change compared with static Young. Data are presented as mean +/- st. deviation, n ≥ 3 per group, in triplicate. * = p<0.05, ** = p<0.01, *** = p<0.001, **** = p<0.0001 as indicated.

Mechanical stretched significantly increased MIP-1β (CCL4) gene expression after 24 hours in Young ATII cells (Fig. 4C). Age significantly increased MIP-1β gene expression, regardless of mechanical stretch condition (Fig. 4 C). The administration of 4-PBA attenuated the increased MIP-1β expression in static or stretched old cells; however, there were no significant differences seen in static or stretched Young ATII cells. Concurrent with the gene expression data for 4 (Fig. 1) and 24 hours (Fig. 4C), MIP-1β cytokine concentration in the media was elevated after 24 hours in stretched and static Old ATII cells compared to stretched and static Young ATII cells (Fig. 4D). Additionally, the administration of 4-PBA decreased both MIP-1β concentrations in all groups (Fig. 4D).

In order to determine the ability of age and stretch to influence ATII recruitment of monocytes, we performed conditioned media experiments by exposing primary BMDMs to ATII conditioned media (CM) from all groups. We first quantified Young and Old BMDM migration using age-matched CM from the ATII stretch experiments (Fig. 5). We observed significantly decreased (p<0.05) BMDM migration with old BMDMs/old static ATII CM in comparison to young BMDMs/young ATII static CM. We also observed this same significant decrease (p<0.05) in migration with old BMDMs/old stretch CM in comparison to young BMDMs/young ATII stretch CM. There was also a significant reduction in migration with young BMDMs/young ATII static CM + 4-PBA in comparison to young BMDMs/young ATII static CM. This same reduction in migration with 4-PBA was also observed in young BMDMs/young ATII stretch CM in comparison to young BMDMs/young ATII stretch CM. Although not statistically significant, 4-PBA appeared to decrease both Young and Old BMDM migration in response to Old ATII CM (Fig. 5).

**Figure 5:**
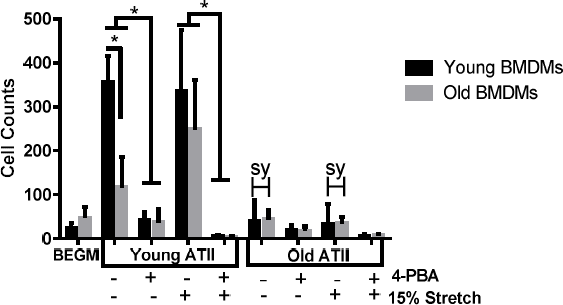
Monocyte Migration with Age-Matched and Mis-Matched ATII Conditioned Media. BEGM is ATII growth media. Data are presented as mean +/- st. deviation, n ≥ 3 per group*p<0.01, “sy”h=statistically significant with p<0.05 compared with Young ATII counterpart. Additional statistically significant differences are described in the Results section.

In order to determine if it was the age of the ATII cells producing the CM or the age of the BMDMs that influenced migration, we quantified migration of young BMDMs with CM from the old stretch and static groups and old BMDMs with conditioned media from the young stretch and static groups to represent Mis-matched CM monocyte recruitment (Fig. 5, gray bars in Young ATII and black bars in Old ATII). Old BMDMs/young ATII stretch CM significantly increased migration, p<0.05, compared with to young BMDMs/old ATII stretch CM. 4-PBA significantly decreased migration in the Old BMDMs/young ATII stretch groups (Fig. 5). Notably, Old ATII CM with Young and Old BMDM caused a significant decrease in migration compared with Young ATII CM in both stretched and static groups (as indicated by “sy”, Fig. 5).

## Discussion

VILI is characterized by an influx of inflammatory cytokines, loss of alveolar barrier integrity with pulmonary edema formation, and altered tissue mechanics. Mechanical ventilation causes alveolar overdistension and other types of lung injury in inflamed regions of the lung (56). The over-distension of aerated lung regions generates abnormally large mechanical stretching forces on the epithelium that causes an immense secretion of proinflammatory cytokines and increased permeability (11, 33). Implementing “protective ventilator strategies” has only marginally improved negative outcomes, and the overall mortality rates for ventilated patients are still unacceptably high (38, 50). Furthermore, few studies are performed on aged subjects, which is incongruent with the fact that elderly patients have a greater need for mechanical ventilation (3,18). These observations illustrate the major clinical need to develop treatments or therapies that prevent the cellular injury mechanisms and inflammation directly resulting from the pathological mechanical forces generated during mechanical ventilations.

We began our study screening for a number of inflammatory cytokines with gene expression. This screening identified a number of genes age-responsive and stretch-responsive in ATII cells. In statically culture, we observed a significant increase in the expression of MCP-1(CCL2), MIP-1α (CCL3), MCP-3 (CCL7), MIP-3α (CCL12), and IL-6st with age. These genes are associated with cytokines and chemokines that result in an increased level of inflammation in the lung (26, 39, 41). MCP-1, MIP-1α, and MCP-5 all showed increased gene expression with age alone, mechanical stretch alone, and with the combination of age and stretch. MCP-1 is a well-studied chemokine that is believed to assist in the recruitment of monocytes, memory T cells, and dendritic cells to sites of inflammation in response to tissue injury or infection (65). It has also been shown to increase in aged mice in response to injury or infection (52). Other studies have shown increased MCP-1, MIP-1a, and IL-6 in experimental VILI models (43). Additionally, MCP-1, MIP-1α, MIP-3α, and IL-6 increase in human epithelial cell and fibroblast cellular senescence (22). Furthermore, we observed increased MIP-1β (macrophage inflammatory protein-1β) and MIP-3α (macrophage inflammatory protein-3α) expression in response to mechanical stretch. MIP-1β is believed to be a chemoattractant for monocytes, natural killer cells, and other immune cell participants (5, 59). While all of these cytokines and chemokines likely play a role in VILI, we selected MCP-1 and MIP-1β as the targets for further study in age-associated ER stress and inflammatory experiments. In addition to its role in monocyte/macrophage recruitment (52), numerous studies indicate that MCP-1 is significantly increased in experimental models of mechanically ventilated mice (27, 28, 30), cyclically stretched murine and human endothelial cells (9, 48, 60), and in lipopolysaccharide (LPS)- induced acute lung injury models using lung epithelial cells (32, 53, 64). While MIP-1β also plays a role in monocyte/macrophage recruitment, studies demonstrate that it is also increased in experimental acute lung injury models (5, 25).

We investigated ER stress as a mediator of the stretch and age-induced inflammation because ER stress plays a significant contributory role in age-related diseases and chronic inflammation (6, 23). Conditions such as hypoxia, calcium ion depletion, oxidative injury, infections, and inflammatory cytokines have the ability to disrupt the ER and prevent the normal protein folding (4, 7). The accumulation of folded and misfolded proteins in the ER leads to ER stress. Failure of the cell to mitigate protein accumulation leads to inflammatory pathway activation (37). Age and/or mechanical stretch likely further disrupt the epithelial cells’ ability to alleviate the unfolded and misfolded protein accumulation in the ER, which leads to intensified inflammation and apoptosis that contribute to barrier dysfunction and increased permeability. While it has been recently shown that ER stress regulates alveolar epithelial homeostasis in response to mechanical stimuli (16), our current study provides the first evidence that aging significantly impacts the ER stress responses and inflammatory signaling in alveolar epithelial cells with and without the addition of mechanical stretch.

In order to attenuate the age-associated increases in mechanical stretch-induced ER stress and inflammation, we administered 4-PBA or vehicle to the ATII cultures. 4-PBA acts as a molecular chaperone to aid in the prevention of misfolded protein accumulation (35). 4-PBA is commercially available, approved by the FDA, clinically used to cure urea cyclic disorders (42). Additionally, several studies have already demonstrated that 4-PBA can be effective in alleviating chronic inflammation and age-related disease outcomes, such as lipopolysaccharide (LPS)-induced lung inflammation in a murine model, through mitigation of ER stress (34, 42). Based upon these prior results, we tested 4-PBA in our *in vitro* cell stretch model.

4-PBA induced changes in concentrations of inflammatory cytokines secreted into the cell media. MCP-1 (CCL2) and MIP-1β (CCL4) media concentrations were significantly decreased with the administration of 4-PBA in both stretched and statically cultured old ATII cells. 4-PBA also mitigated MCP-1 secretion in young static and stretched cells, while MIP-1β secretion did not change in the young static or stretched cells. The reduction in these concentrations may also be the cause for the significantly reduced monocyte recruitment as well, as they are believed to help regulate monocyte and other immune cell recruitment and activation (5, 52, 54).

Macrophages are thought to be one of the major targets of the mechanically stretch-induced signaling mechanisms of the alveolar epithelium (55, 58). Macrophages in the alveolar space greatly contribute to barrier integrity and local inflammation as they are a major participant in inflammatory signaling between the epithelium and innate immune response (8, 10). We investigated the impact of age and/or stretch on monocyte recruitment using BMDMs to understand the interaction between macrophages and epithelial cell signaling. We chose to use this primary cell type in order to examine age-related differences. Large quantities are more easily obtained with BMDMs compared with alveolar macrophages, and they are more suitable for microscopy applications (57). Based on our age-matched data alone, we were unable to determine if it was the age of the BMDMs or the age of the epithelial cells that elicited the significantly different responses. Our age mismatched experiments (Fig. 5) suggest that age of the ATII cells may have a greater impact on BMDM recruitment than the age of the BMDMs themselves. Administration of 4-PBA significantly reduced BMDM recruitment with young BMDMs/young static CM, young BMDMs/young stretch CM, and with old BMDMs/young stretch CM. These results indicate that 4-PBA may quell inflammation and macrophage recruitment in stretch injury.

There are some additional limitations in this the study. Multiple studies have suggested that 15% stretch is insufficient to injure young alveolar epithelial cells (11, 15). This possibility might explain why we did not see the same inflammatory or ER stress response changes that we observed in the old ATII cells. However, our results suggest that advanced age impacts inflammatory and ER stress activation in ATII cells in response to mechanical stimuli. We have shown previously that while cell membrane integrity is retained, cyclic stretch of young ATII cells at 15% change in surface area is sufficient to affect gene expression and phenotype (29). Our results indicate that old ATII cells respond differently to mechanical stretch compared with young ATII cells, potentially indicating that even under low tidal volume mechanical ventilation, older subjects may have an altered inflammatory response.

As the compounding effects of aging on lung injury and inflammation are becoming increasingly recognized, these age-dependent factors that may be associated with the injury and inflammation responses between the alveolar epithelium and innate immune system still need more clarity. While the administration of 4-PBA shows promise in treating the ER stress and inflammation responses following mechanical stretch, the age‐ and stretch-dependent mechanisms and the use of 4-PBA to mitigate inflammation and ER stress require further study to prove their practicality for future lung injury-related clinical potential.

## Acknowledgements

The authors acknowledge the assistance of Niraja Bohidar in aiding in cell isolation experiments. This research was supported by the National Institutes of Health under Award Number R01AG041823. The content is solely the responsibility of the authors and does not necessarily represent the official views of the National Institutes of Health.

